# lncRNAKB: A comprehensive knowledgebase of long non-coding RNAs

**DOI:** 10.1101/669994

**Authors:** Fayaz Seifuddin, Komudi Singh, Abhilash Suresh, Yun-Ching Chen, Vijender Chaitankar, Ilker Tunc, Xiangbo Ruan, Ping Li, Yi Chen, Haiming Cao, Richard S. Lee, Fernando Goes, Peter P. Zandi, M. Saleet Jafri, Mehdi Pirooznia

**Author notes:** To whom correspondence should be addressed to: Mehdi Pirooznia, MD., PhD., Director, Bioinformatics and Computational Biology, National Heart Lung and Blood Institute, National Institutes of Health, https://bioinformatics.nhlbi.nih.gov, BG 12A RM 3053N, 12 SOUTH DR., Bethesda MD 20892.

## Abstract

We have assembled a comprehensive <u>l</u>ong <u>n</u>on-<u>c</u>oding <u>RNA</u> <u>k</u>nowledge<u>b</u>ase (lncRNAKB) of 77,199 annotated human lncRNAs (224,286 transcripts) by methodically integrating widely used lncRNAs resources. To facilitate functional characterization of lncRNAs, we employed Genotype-Tissue Expression (GTEx) project to provide tissue-specific gene expression profiles of lncRNAs in 31 solid organ tissues. Additional information includes network analysis to identify co-expressed gene modules to potentially delineate lncRNA function. Tissue-specificity, phylogenetic conservation scores and coding potential for lncRNAs are included. Finally, using whole genome sequencing data from GTEx, expression quantitative trait loci (*cis*-eQTL) regulated lncRNAs were calculated in all tissues. lncRNAKB is available at http://www.lncrnakb.org.

## BACKGROUND

A majority of the human genome is non-protein-coding and was considered “junk DNA” when scientists of the Human Genome Project presented the first rough draft of the human genome sequence [1,2]. However, that notion is drastically changing with the introduction of high throughput technologies such as Next-Generation Sequencing (NGS) that have allowed the non-coding genome to be interrogated at high resolution and scale [3]. The Encyclopedia of DNA Elements (ENCODE) project reports that only ~2% of the genome is protein-coding; however, approximately 80% of the non-protein coding genome are detectably transcribed under some conditions [4]. Among the non-protein coding transcripts, the long non-coding RNAs (lncRNAs) are a class of transcripts whose length range from 200 base pairs (bp) to 100 kilobases (kb) (approximately 10 kb on average) [5]. Currently, the number of estimated lncRNAs annotations in humans range from 20,000 to 100,000 [6].

While the field of lncRNA is nascent, enough evidence exist to suggest that they play a critical role in many biological processes including transcriptional and post-transcriptional regulation, epigenetic regulation, organ or tissue development, cell differentiation and apoptosis, cell cycle control, cellular transport, metabolic processes and chromosome dynamics [7,8]. A consensus has emerged on other features of lncRNAs such as lack of interspecies conservation [9–12], low level of expression [7], and higher degree of tissue-specific expression compared to mRNAs [10,13]. The little sequence conservation among lncRNAs is confined to short, 5′-biased patches of conserved sequences nested in exons [9]. Some lncRNAs undergo translation, though only a minority of such translation events results in stable and functional peptides [14,15].

Recently, several publicly available resources dedicated to annotation of lncRNAs in humans and other species have been developed [6,16,16,17]. Most of these databases are available through web-based searchable interfaces and provide downloadable annotation files in Gene Transfer Format (GTF) or Gene Feature Format (GFF) [18–20]. Frequently used resources of lncRNAs annotation include GENCODE [21,22], CHESS [23], LNCipedia [24,25], NONCODE [26], FANTOM [27], MiTranscriptome [28] and BIGTranscriptome [3]. These resources annotated lncRNAs by two approaches: manual or automatic [6]. Manual annotation involves human annotators curating gene and transcript models based on RNA and protein experimental evidence and defined sets of rules [22]. Automatic annotation uses bioinformatics methods such as StringTie [29] and Cufflinks [30] to reconstruct gene and transcript models based on billions of short RNA-sequence (RNA-seq) reads [28]. A few of these databases have attempted to integrate annotations from multiple sources such as expression, methylation, variation, conservation and functional annotation of lncRNAs in humans. The information on lncRNAs provided in these databases is, however, not comprehensive and at times inadequate, making it difficult to understand their molecular and cellular functions.

To address the limitations and shortcomings of the existing lncRNA databases, we have developed lncRNAKB which systematically combines the frequently used lncRNAs annotation resources mentioned above using a cumulative stepwise intersection method. This method of integration compares the annotations thoroughly, discarding redundant/ambiguous lncRNAs records and also accounts for the large overlap between the lncRNAs annotation databases to avoid redundancy. The lncRNAKB also provides a comprehensive downloadable, searchable and viewable (via the UCSC Genome Browser) [31] GTF annotation file of human PCGs and a large number of lncRNAs (*n*=77,199) that can be used by researchers to quantify RNA-seq data generated in lab or downloaded from public domains for lncRNA discovery. To improve lncRNAs’ functional characterization, we have implemented an up-to-date analysis pipeline to analyze the tissue-specific RNA-Seq data available through Genotype Tissue Expression (GTEx) project [32] and have created a tissue-specific expression body map of human lncRNAs. Using the gene expression information, we calculated tissue-specificity scores across all the genes in the lncRNAKB. Additionally, we calculated expression quantitative trait Loci (eQTL)-regulated lncRNA genes using the GTEx expression and GTEx whole genome sequencing (WGS) genotype data and created a tissue-specific eQTL body map of human lncRNAs. Additional features of the lncRNAKB include providing information regarding classification of lncRNA based on their positional information and calculating their coding potential using FlExible Extraction of LncRNAs (FEELnc) [33] algorithm. Furthermore, the knowledgebase also provides eQTL results and exon-level conservation scores derived from an alignment of 30 vertebrate species [31] for all PCGs and lncRNAs. Finally, to predict lncRNA functions, we used Weighted Gene Co-expression Network Analysis (WGCNA) [34] method to analyze lncRNAs-mRNAs co-expression patterns in a tissue-specific manner. The co-expression modules were subjected to pathway enrichment analysis to identify functional pathways associated with lncRNAs thereby creating a tissue-specific body map of functionally annotated lncRNAs. All pathway results files can be browsed or are available for download. Moreover, for each tissue we have selected 25 notable pathways and created a dynamic network figure on the website to view the strength of connections between the highest-ranking mRNAs and lncRNAs. All these features in combination with annotation of 77,199 lncRNAs presented in the lncRNAKB provides the most up-to-date and comprehensive functional information about lncRNAs.

## CONSTRUCTION and CONTENT of lncRNAKB

### Data sources and collection

To identify widely used lncRNAs databases we performed literature search of the PubMed database through February 28^th^, 2019 with the following keyword algorithm: *(lncrna or long noncoding or long non-coding rna or noncoding) and (annotation or function or database).* A total of 13,412 articles were returned filtered by human species and published within the past five years sorted by the best match criteria. The titles, abstracts, keywords, and full text were manually reviewed to identify publications that reported lncRNAs annotations, databases and function. The references of these articles were also searched to identify other articles that were potentially missed by the initial PubMed search. After this review, six lncRNA resources were selected for step-wise integration to create lncRNAKB. The six resources are: CHESS (version 2.1), LNCipedia (v5.2), NONCODE (v5.0), FANTOM (5.0.v3), MiTranscriptome (v2) and BIGTranscriptome (v1).

### Data integration

The gene transfer format (GTF) or gene feature format (GFF) from all six annotation databases (links in Table 1) were downloaded. To streamline the data integration step, all the GTF or GFF annotations were parsed to the same format using the following steps: (i) where required, the coordinates of annotation were updated using the UCSC liftOver tool [31] from hg19 to hg38 (latest genome build), and (ii) for each chromosome, the gene and transcript records were split into individual files named by chromosome, strand, start and stop base pair locations. Each gene block file contained the transcripts information and the transcript block file contained the exons information. In cases where the transcripts or exons records lacked genes information, a gene entry was manually created using the gene ids in the transcripts or exons records and combined with the base pair locations of the first exon (as gene start), of the last exon (as gene stop), and transcript strand to represent the gene strand. All the redundant records between annotation files were removed in this process.

**Table 1:** Summary of lncRNAs annotation databases that have been integrated into the lncRNAKB and the types of data included in each resource *lncRNAs only

|  | CHESS2.1 | LNCipedia5.2 | NONCODEv5.0 | FANTOM5.0.v3 | MiTranscriptomev2 | BIGTranscriptomev1 | lncRNAKB |
| --- | --- | --- | --- | --- | --- | --- | --- |
| Annotation source file name | chess2.1.gff | lncipedia_5_2_hc_hg38.gtf | NONCODEv5_human_hg38_lncRNA.gtf | FANTOM_CAT.lv3_robust.only_lncRNA.gtf | mitranscriptome.hg19.v2.gtf | BIGTranscriptome_lncRNA_catalog.hg19.gtf | lncRNAKB_hg38_v6.gtf |
| Website | http://ccb.jhu.edu/chess/ | https://lncipedia.org/info | http://www.noncode.org/index.php | http://fantom.gsc.riken.jp/5/ | http://mitranscriptome.org/ | http://bhyou.dothome.co.kr/ | https://www.lncrnakb.org |
| Reference[PMID] | Pertea *et al.* , 2018 [30486838] | Volders *et al.* , 2013, 2015, 2019 [30371849] | Fang *et al.* , 2018 [29140524] | Hon *et al.* , 2017 [28241135] | Iyer *et al.* , 2015 [25599403] | You *et al.* , 2017 [28396519] | Seifuddin *et al.* |
| Genome reference build/version | hg38 | hg19, hg38 | hg19, hg38 | hg19 | hg19 | hg19 | hg38 |
| Annotation method/source | GENCODE (release 25 and 27), RefSeq, FANTOM5, RNA-seq (GTEx-phs000424.v6.p1 in May of 2016), transcript assembly, mass spectrometry (validation) | Ensembl, RNA-seq (Human BodyMap 2.0 lincRNAs), lncRNadb, GENCODE (release 13), RefSeq-Dec2014, RefSeq-NCBI (release 106), Nielsen *et al.*, Hangauer *et al.*, NONCODE, Sun and Gadad *et al.*, 2015, FANTOM CAT | Ensembl, RefSeq, lncRNadb, LNCipedia, RNA-seq (Human BodyMap 2.0 lincRNAs), Exosome Expression Profile, old versions of NONCODE | GENCODE (release 19), Human BodyMap 2.0, miTranscriptome, ENCODE and an RNA-seq assembly from 70 FANTOM5 samples, cap analysis of gene expression (CAGE) data | 7,256 RNA sequencing (RNA-seq) libraries from tumors, normal tissues and cell lines comprising over 43 Tb of sequence from 25 independent studies including ENCODE, TCGA, Human BodyMap 2.0, proteomics (validation) | ENCODE, TCGA, GTEx, Human BodyMap 2.0, the Human Protein Atlas, GENCODE (release 19), RefSeq, PacBio | CHESS2.1, LNCipedia5.2, NONCODEv5.0, FANTOM5.0.V3, MiTranscriptomev2, BIGTranscriptomev1 |
| Number of lncRNAs (genes) | 19,897 | 56,946 | 96,308 | 27,871 | 63,505 | 14,090 | 77,199 |
| Number of lncRNAs (transcripts) | 56,922 | 127,802 | 172,216 | 89,833 | 175,259 | 26,591 | 224,286 |
| Number of lncRNAs (exons) | 205,217 | 357,620 | 429,240 | 251,201 | 539,840 | 87,316 | 611,340 |
| Number of protein-coding genes | 22,883 | * | * | * | * | * | 22,518 |
| Tissue-specific Expression/score | yes | no | yes | yes | yes | yes | yes |
| Tissue-specific Expression Quantitativ Trait Loci (eQTLs) | no | no | no | no | no | no | yes |
| UCSC genome browser/Custom Genome Browser Track | no | yes | yes | yes | yes | no | yes |
| External Gene information Links (Gene Cards or RefSeq or Ensembl or UCSC) | yes | yes | yes | yes | yes | yes | yes |
| Coding-potential prediction/score | no | yes | no | yes | yes | yes | yes |
| Conservation information | no | yes | yes | yes | yes | no | yes |
| Gene-level functional annotation, mRNA coexpression, Pathway enrichment analysis | no | no | yes | yes | yes | no | yes |

Using CHESS as the reference annotation database (containing both protein-coding and lncRNAs genes) we used a cumulative stepwise intersection method to merge it with the rest of the five lncRNAs annotation databases in the following order: (i) FANTOM, (ii) LNCipedia, (iii) NONCODE, (iv) MiTranscriptome and (v) BIGTranscriptome at the genes and transcripts levels. This order of intersection was arbitrarily chosen. Figure 1 illustrates the cumulative stepwise intersection method for two databases as an example, D1 (CHESS) in blue and D2 (FANTOM-lncRNAs only) in green. For each gene entry in D1 (top blue panel), we kept genes from D2 (green panel) that had full overlap and were also within D1 gene boundary. The resulting intersection is shown in orange. D2 gene that had partial overlap with D1 gene (marked in red X) were discarded as we did not want to re-define boundaries of genes in the reference annotation database.

**Figure 1:**
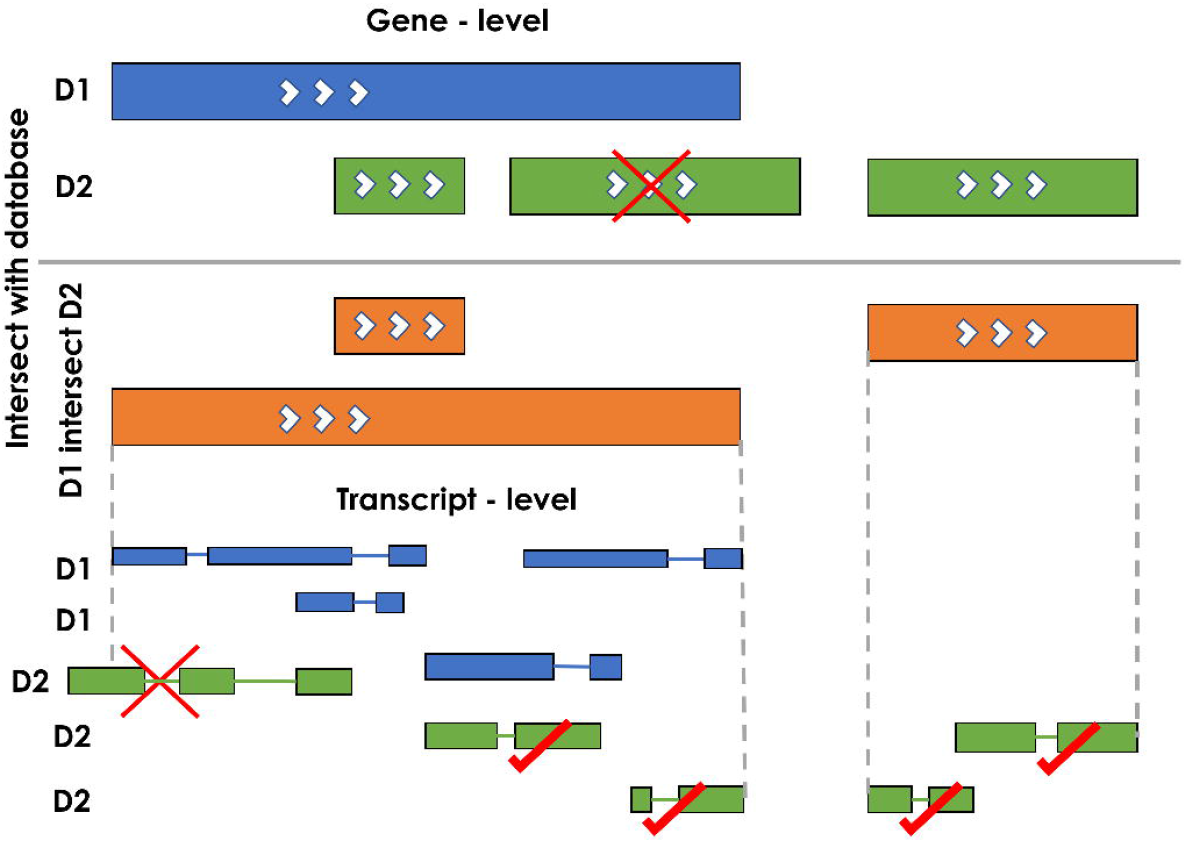
Illustration showing the stepwise intersection of two annotation databases D1 (CHESS) (blue) and D2 (FANTOM-lncRNAs only) (green) at the gene and transcript levels. The genes are shown as solid rectangles and the transcripts are shown with exons and introns. The white arrows show the direction/strand in which the gene is transcribed. The orange bars show the results of the intersection (D1 intersect D2) at the gene level. The red X marks show transcripts that were not incorporated into the merged annotation and vice versa for the red ticks. D3 (LNCipedia), D4 (NONCODE), D5 (MiTranscriptome) and D6 (BIGTranscriptome) were merged using the same cumulative stepwise intersection method (see construction and content of lncRNAKB)

For genes that intersected, the transcript records (shown as smaller bar connected by line to represent exons and introns, respectively) from D1 and D2 were compared. Similarly, to the gene intersection, transcript entries whose start and stop were within the gene boundaries were included. Several transcripts (marked with a red X) that fell outside the gene boundary and were probably incorrectly assigned to genes were removed in this process. In addition, if a transcript in D2 had partial or no overlap with transcripts in D1, we incorporated that transcript (marked with red checks) including all the exons to the gene record accordingly. For genes with no overlap in D1, we added all the transcripts and corresponding exons to the merged annotation as a lncRNA entry (marked in red checks).

### Evaluation of coding potential of lncRNAs

FlExible Extraction of LncRNAs (FEELnc) [33] was used to classify/annotate and calculate the coding potential of all the gene entries in the lncRNAKB. FEELnc annotates lncRNAs based on a machine learning method, Random Forest (RF) [35], trained with general features such as multi *k*-mer frequencies, RNA sequence length and open reading frames (ORFs) size. It is comprised of three modules: (i) filter, (ii) coding potential, and (iii) classifier. The filter module flags and removes transcripts overlapping (in sense) exons of the reference annotation, specifically the protein-coding exons. We used the GENCODEv29 [22] GTF file as the reference annotation to get an estimate of the number of transcripts from lncRNAKB overlapping with “*protein_coding*” transcripts. We arbitrarily set the minimal fraction out of the candidate lncRNAs size to be considered for overlap to be excluded as 0.75 (> 75% overlap) to retain many lncRNAs transcripts. Transcripts < 200 base pairs (bp) long were filtered out and monoexonic transcripts were retained. We then used the filtered GTF annotation output file from the filter module and calculated a coding potential score (CPS) for each transcript using the coding potential module. Due to the lack of a gold standard/known human lncRNAs data set for training, we used the “intergenic” mode in the module. This approach extracts random intergenic sequences of length *L* from the genome of interest to model species-specific noncoding sequences as the non-coding training set. We used the human reference genome FASTA file (hg38) and the GENCODE GTF file as the reference annotation. To get the best training set of known mRNAs, we used *“transcript_biotype=protein_coding*” and “*transcript_status=KNOWN”* for the RF model. We used the default values for the *k*-mer sizes, number of trees and ORF type. To determine an optimal CPS cut-off, FEELnc automatically extracts the CPS that maximizes both sensitivity and specificity based on a 10-fold cross-validation. The CPS was between 0 and 1 where 0 indicates a non-coding RNA and a score close to 1 a mRNA. And finally, to classify potential lncRNAs with respect to the localization and the direction of transcription of nearby mRNAs (or other non-coding RNAs) transcripts as shown in Supplementary Figure 1, we used the classifier module. We used the final set of lncRNAs transcripts output from the coding potential module and classified them using the GENCODEv29 GTF file as the reference annotation. A sliding window size around each lncRNA was used to check for possible overlap with nearest reference transcripts. We used a minimum and maximum window size of 10 kilobase (kb) and 100kb respectively. The classification method reported all interactions within the defined window and established a best partner transcript using certain rules.

### Conservation Analysis

Conservation of exons between protein-coding genes and lncRNAs in the lncRNAKB annotation database was analyzed using the bigWigAverageOverBed [36] and the cons30way (hg38) track [37] both downloaded from the UCSC genome browser. This track shows multiple alignments of 30 vertebrate species and measurements of evolutionary conservation using two methods (phastCons and phyloP [38]) from the PHAST package [39] for all thirty species. The multiple alignments were generated using multiz [40] and other tools in the UCSC/Penn State Bioinformatics comparative genomics alignment pipeline. An exon-level BED file was created using the lncRNAKB GTF annotation file separately for protein-coding genes and lncRNAs. We merged overlapping exons within transcripts to avoid counting conservation scores of overlapping base pairs more than once. For each exon, the bigWigAverageOverBed function calculates the average conservation score across all base pairs. Using boxplots, we visualized and compared the average conservation score differences between lncRNAs and protein-coding exons.

### Expression profiling

To evaluate gene expression, RNA-seq data available through the Genotype Tissue Expression (GTEx) project [32] was utilized. The raw paired-end RNA-seq data (FASTQ files – GTEx Release v7) from the dbGap portal (study_id=phs000424.v7.p2) of 31 solid organ human normal tissues were downloaded. For each solid tissue, quality control of paired-end reads were assessed using FastQC tools [41], adapter sequences and low-quality bases were trimmed using Trimmomatic [42] and aligned to the human reference genome (*H. sapiens*, GRCh38) using the HISAT2 [43]. Using the uniquely aligned reads to the human genome, gene-level expression (raw read counts) were generated with the featureCounts software [44] using the lncRNAKB GTF annotation. After visualizing the distribution of uniquely mapped paired-end reads assigned to genes across all the GTEx samples, samples with < 10^6^ reads assigned to genes were excluded. In addition, there were samples with data that we could not map or download. We normalized the raw read counts to Transcripts Per Kilobase Million (TPM) [45]. To explore gene expression similarity between tissues and across GTEx samples as well as summarize lncRNAs tissue-specific expression we performed a principal component analysis (PCA) using the prcomp package in R [46,47]. We used the normalized TPM expression values, transformed by taking the *log*_2_(*TPM*), across all lncRNAs (*n* = 77,199) and tissues (*n* = 31) (no filters applied).

### Tissue-specificity scores

In addition to gene expression quantitation, we calculated two tissue-specificity metrics (Tau and Preferential Expression Measure (PEM)) [48,49] using the normalized TPM expression values across all genes and tissues (no filter applied). Tau summarizes in a single number whether a gene is tissue-specific or ubiquitously expressed across all tissues. PEM shows for each tissue separately how specific the gene is to that tissue. The PEM scores the expression of a gene in a given tissue in relation to its average expression across all other genes and tissues. The average gene expression across all replicates by tissue was used to compute Tau and PEM. Genes that were not expressed in at least one tissue were removed from the analysis.

### Genotype file processing

The whole genome sequence (WGS) data in blood-derived DNA samples from GTEx portal (dbGaP: phs000424.v7.p2) was downloaded to conduct tissue-specific expression quantitative trait loci (eQTL) analysis. First, the VCF files were processed using the following steps with a combination of PLINKv1.9 [50,51] vcfv0.1.15 [52] and bcfv1.9 tools [53]: (i) remove indels; (ii) exclude missing and multi-allelic variants; (iii) selected “FILTER == ‘PASS’” variants; (iv) exclude variants with minor allele frequency (MAF) < 5%; (v) update the coordinates of single nucleotide polymorphisms (SNPs) using the UCSC liftOver tool [31] from hg19 to hg38 (latest genome build); (vi) change the SNPs IDs to dbSNP [54] rsID using dbSNP Build 151; (vii) convert to bed, bim and fam format. For each solid tissue, subjects with both WGS data and gene expression data were selected. The VCF file was subset by tissue and the MAF recalculated to exclude variants with MAF < 5%. After converting to ped and map format, we ran principal component analysis (PCA) on each tissue to get a set of genotype covariates using eigensoftv6.1.4 [46,55].

### eQTL analysis

For each solid tissue, we implemented a two-step filtering approach, which is similar to the steps adapted by GTEx [32]. Briefly, the genes were first filtered based on TPM to include genes with TPM > 0.50 in at least 20% of the samples. Next, the genes were filtered based on raw counts to include genes with counts > 2 in at least 20% of the samples. The edgeR [56] and limma-voom [57] [58] package in R [59] were used to process the filtered read counts into log2 counts per million (log2CPM) that were normalized using trimmed mean of M-values (TMM) [60]. The expression files were then sorted by gene start and stop, compressed with BGZIP and indexed with TABIX [61]. Only tissues with > 80 samples were included in the *cis*-eQTL analysis. For eQTL analysis, the first five principal components (PCs) (see Genotype file processing) and sex information was included as a covariate. Within each tissue, *cis*-eQTLs were identified by linear regression, as implemented in FastQTLv2.0 (threaded option) [62], adjusting for the five PCs and sex. We restricted our search to variants within 1 megabase (Mb) of the transcription start site (TSS) of each gene. In addition, the adaptive permutations option in FastQTL between 1000 and 10000 permutations. Once we obtained the permutation p-values for all the genes, we accounted for multiple testing to determine the significant *cis*-eQTLs. We used the Benjamini and Hochberg correction method [63] to calculate the false discovery rate (FDR) in R statistical programming language (R) [59]. For each tissue, all *cis*-eQTL results were visualized using a Manhattan plot created using the qqman package in R [64].

### Functional characterization of lncRNAs using a network-based approach

Using the filtered log2CPM and TMM normalized gene expression data (see construction and content of lncRNAKB: Expression profiling and eQTL analysis), we used the weighted gene co-expression network analysis (WGCNA) approach [34] as implemented in the Co-Expression Modules identification Tool (CEMiTool) package in R [65] to identify modules of lncRNA-mRNA clusters that are co-expressed and therefore likely work in concert to carry out various biological functions. For this, the gene expression data was filtered by log2CPM > 2 in at least 50% of the samples to avoid random correlations between low-expressing genes. The default CEMiTool parameters were used with the following exceptions: (i) Pearson method was used for calculating the correlation coefficients, (ii) the network type used was unsigned, (iii) no filter was used for the expression data, (iv) applied Variance Stabilizing Transformation (VST) and the correlation threshold for merging similar modules were set to 0.90. All the co-expressed modules were subjected to over-representation analysis (ORA) by module based on the hypergeometric test [66]. We used Gene Ontology (GO) pathways [67–69] to check for overrepresentation of genes and determined the most significant module functions based on pathways FDR q-value ≤ 0.05 [70]. The background set used for the pathway enrichment analysis was genes represented across all GO pathways. To visualize the interactions between the genes in each co-expression module, we selected 25 notable pathways for each tissue. The module adjacency matrices for each of these pathways were filtered based on correlations > 0.20 across all genes in each pathway. A JSON file (one per pathway) was created to produce interactive networks using Cytoscape v3.6.0 JavaScript modules [71]. The network files and the module adjacency/correlation matrix files are available for downloading at lncRNAKB.

### Architecture of the database

The 3-tier server architecture model containing data, logic and presentation tiers has been implemented as shown in Figure 2. The popular MySQL open source relational database management system (RDBMS) has been employed for the data tier, expanded with a NoSQL document storage. NoSQL document storage is a JSON-based (JavaScript Object Notation) data structure format and as such has a flexible dynamic structure with no schema constraints which makes it suitable for literature and document storage. The MySQL RDBMS is ideal for data indexing and a powerful query system for relational data. The logic tier is responsible for the communication between the user queries from the presentation tier and fetching the outcome from the data tier, as well as data integration from MySQL and NoSQL data sources. The presentation tier contains several modules based on AJAX (Asynchronous JavaScript and XML), jQuery (JavaScript Query system version 3.3.1 - https://jquery.com/), and the PHP server-side scripting language (version 7.1.18.), as well as the CSS (Cascading Style Sheets) code to describe how HTML elements are to be displayed on user side web interface. jQuery and AJAX have the advantage of asynchronous background calls to the logic tier, native JSON parsing, and dynamic rendering of the browser display, which makes the data retrieval system perform more efficiently. The Web server is hosted on a CentOS 7 operating system using an Apache (2.4.33) web server. The user interface is functional across major web-browsers such as Chrome, Safari, and Firefox on Linux, Mac, iOS, Android, and Windows OS platforms.

**Figure 2:**
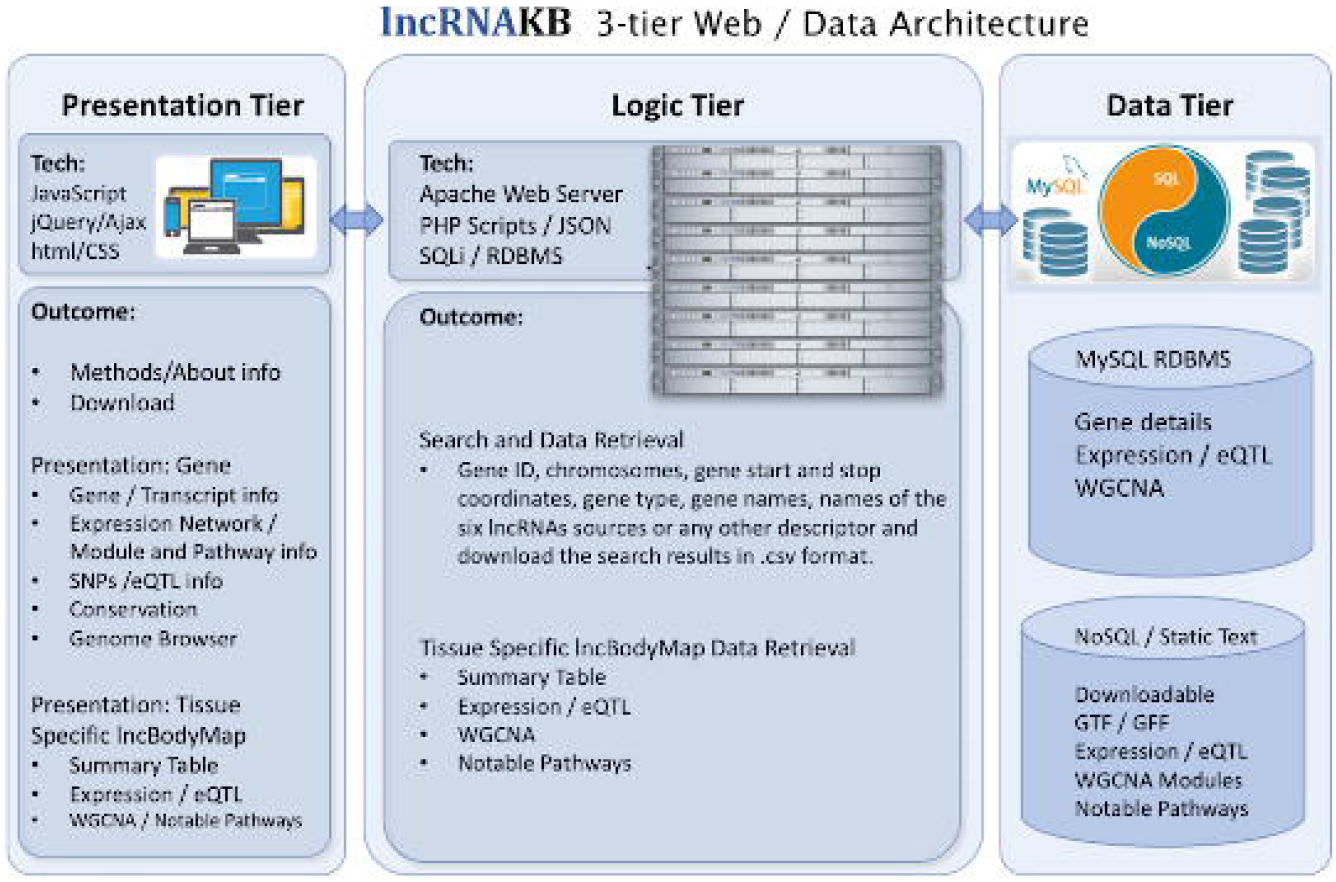
Schema of the web/database segment of the lncRNAKB

## UTILITY and DISCUSSION

### Database content

The lncRNAKB is the end product showcasing the results of a number of analysis that were performed with the goal to provide a **1. comprehensive annotation** and **2. complete functional characterization of lncRNAs**. First, a systematic integration of lncRNA entries from six databases resulted in annotation of 77,199 lncRNAs. Additionally, the integration methodology used also enabled addition of a number of lncRNA transcripts to the known protein-coding gene entries (see construction and content of lncRNAKB). Every gene and/or transcript entry were accompanied by addition of respective exon entries. The resulting annotated lncRNA entries can be browsed or a gene transfer format (GTF) comprising of these annotations is available for download via the Website (see utility and discussion: Downloadable, searchable and viewable lncRNA database). Next, to supplement the lncRNA annotation with functional information a number of additional analysis were performed, and the results can be viewed in the website. In summary, all the lncRNAs were classified based on their position, their coding potential, conservation scores were evaluated (see utility and discussion: Evaluation of coding potential of lncRNAs and Evaluation and comparison of lncRNA and mRNA conservation scores). Additionally, their gene expression across all tissues were evaluated and their tissue specificity score determined (see utility and discussion: Tissue-specific expression profiling of lncRNAs and Evaluating tissue-specificity of lncRNAs). Furthermore, *cis*-expression quantitative trait loci (*cis*-eQTL) regulated lncRNAs were identified (see utility and discussion: eQTL analysis of lncRNAs). Finally, lncRNA-mRNA co-expression networks were evaluated and functionally annotated using pathway enrichment analysis (see utility and discussion: Functional characterization of lncRNAs using a network-based approach). Figure 3 illustrates the results of the lncRNAKB that are discussed in detail in each section below. Table 1 summarizes the number of lncRNAs features (genes *n* = 77,199, transcripts *n* = 224,286 and exons *n* = 611,340) in the lncRNAKB GTF annotation file.

**Figure 3:**
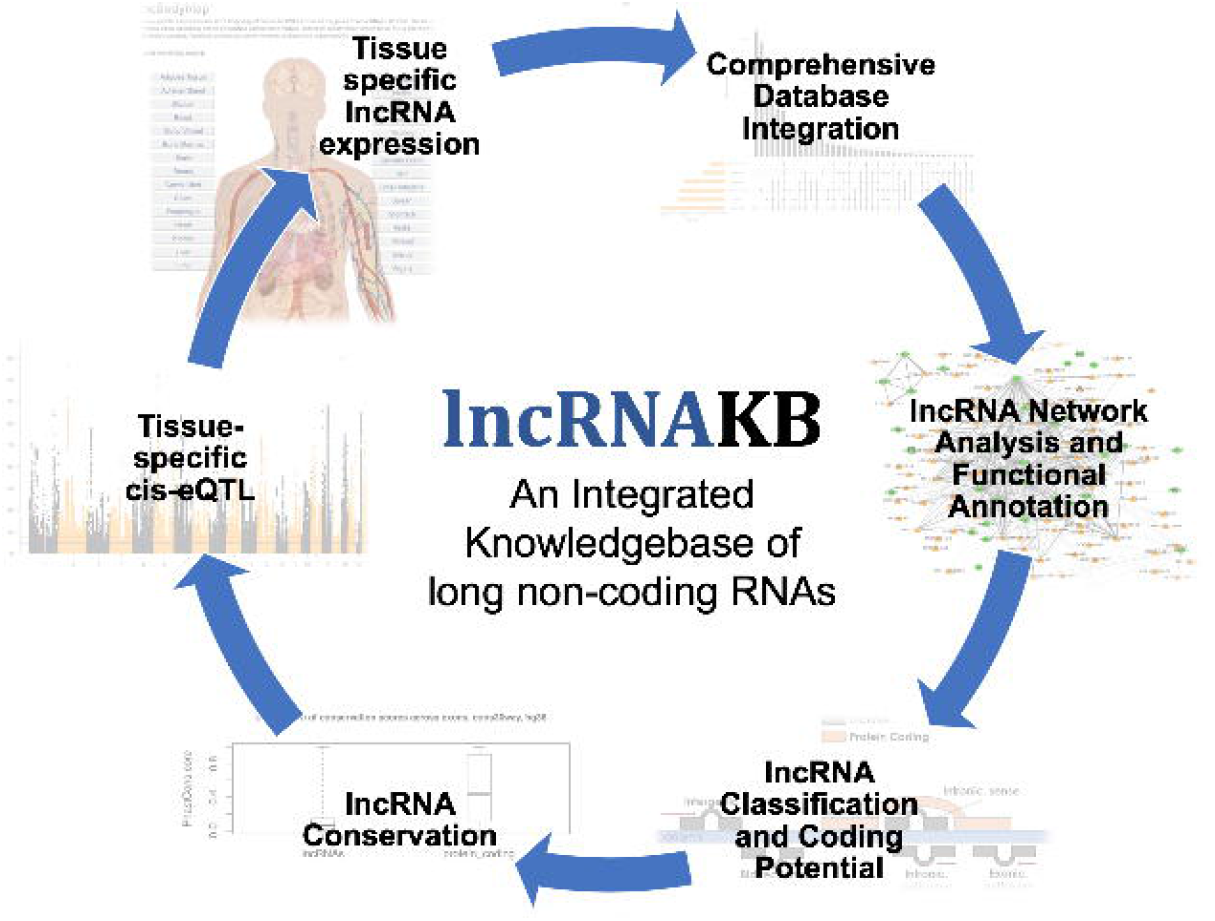
All components of the lncRNAKB which provide valuable information on lncRNAs and are freely available for viewing and downloading on the web resource

### Downloadable, searchable and viewable lncRNA database

Based on the PubMed search and literature review, six databases were chosen to systematically integrate all the lncRNA entries with the goal of providing a comprehensive annotation of lncRNAs (see construction and content of lncRNAKB). Table 2 shows the results of the cumulative stepwise intersection method across the six lncRNAs annotation databases compared to the reference (CHESS) at the gene level. The order in which the six databases are listed in this table is the same order in which the databases were integrated. Briefly, NONCODE and MiTranscriptome added 20,700 and 15,164 genes respectively. While CHESS already incorporated data from FANTOM, based on the cumulative stepwise intersection method we added additional 7,157 genes from FANTOM. LNCipedia on the other hand added 10,506 genes. The last source, BIGTranscriptomev1 contributed only 333 genes which indicates that there was extensive overlap with other annotation databases.

**Table 2:** Results of the cumulative stepwise intersection method across the six lncRNAs annotation databases compared to the reference (CHESS) at the gene level ^1^The original number of genes in the sources shown here are slightly less than the actual downloaded GTF/GFF annotation files (Table 1) because we removed redundant genes and transcripts records (see construction and content of lncRNAKB)

| CHES2.1 | lncRNAKB_hg38_v2.gtf | source | <sup>1</sup>Original number of genes in source after lncRNAKB filter |
| --- | --- | --- | --- |
| 46,421 | 45,857 | CHES2.1 | 46,421 |
|  | 7,157 | FANTOM5.0.v3 | 21,457 |
|  | 10,506 | LNCipedia5.2 | 40,188 |
|  | 20,700 | NONCODEv5.0 | 63,055 |
|  | 15,164 | MiTranscriptomev2 | 45,282 |
|  | 333 | BIGTranscriptomev1 | 13,525 |
| **Total** | **99,717** |  | **229,928** |

Figure 4 illustrates contribution of lncRNAs from each of the six databases. This figure also highlights that there was considerable overlap between different sub-sets of the annotation databases. All of LNCipedia genes overlapped with one or more of the other five annotation databases. NONCODE added the highest number of non-overlapping genes (*n* =16,080) followed by MiTranscriptome (*n* =14,620). BIGTranscriptome added only 333 unique gene entries due to its overlap with genes in the other databases. CHESS was used as the reference annotation database and contains protein-coding (*n* =20,352) and lncRNAs genes (*n* = 18,897). However, from Figure 4, we observed that the number of non-overlapping genes added from CHESS is 9,595, which indicates that we added non-coding transcripts from overlapping lncRNAs in other annotation databases to the protein-coding genes. 5,295 genes overlapped between all six sources. Supplementary Table 1a and 1b shows the number of transcripts and the sources of annotation databases at gene level for protein-coding genes between CHESS and lncRNAKB, respectively. Comparing these two tables showed that the number of transcript entries for the protein coding genes in lncRNAKB was much higher than that in CHESS (approximately 40,330 more transcript entries in lncRNAKB compared to CHESS). This suggests that a good proportion of the lncRNAs transcripts (~15%) overlap with or fall within the boundary of protein coding genes. Supplementary Table 2a and 2b shows the number of transcripts and the sources of annotation databases at gene level for non-coding genes between CHESS and lncRNAKB, respectively. By comparing all 4 tables, we show that we have effectively added numerous non-coding genes (*n* = 77,199) and non-coding transcripts (*n* = 224,286) from different lncRNAs annotation databases.

**Figure 4:**
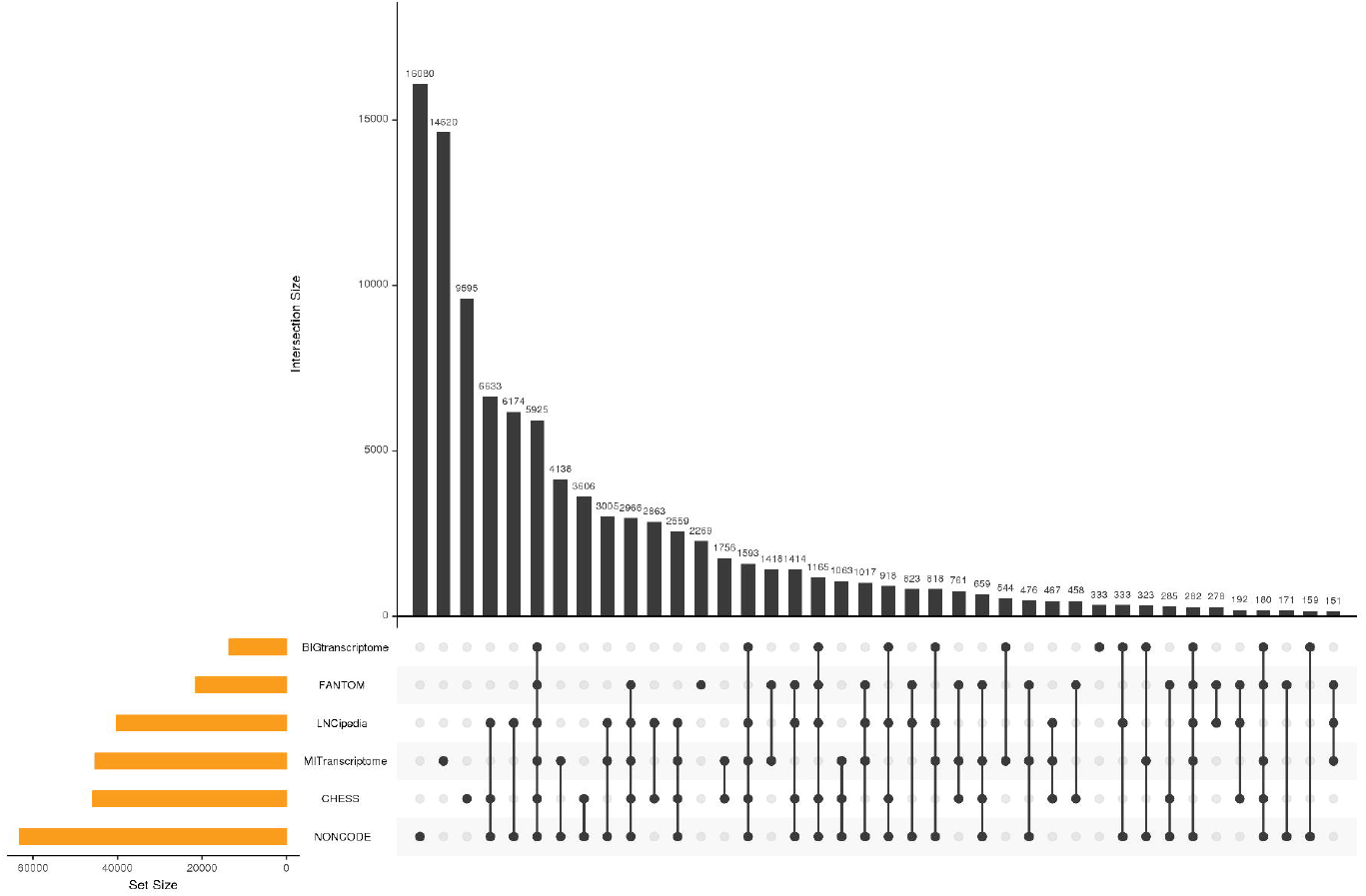
Upset plot showing the overlap of all six lncRNAs annotation databases at the gene level, after the cumulative stepwise intersection method across. The orange bars indicate the total number of genes in each source before merging. The black bars indicate the total number of genes present within a database or shared between databases indicated by black dots present below the x-axis of the plot. Genes uniquely contributed by a single database would be represented as a single dot that horizontally aligns with the respective database. Black dots connected by lines indicate the number of databases that share the genes represented in the bar plot.

To summarize, the final merged annotation in lncRNAKB that comprises both the protein-coding and lncRNAs has 99,717 genes, 530,947 transcripts, 3,513,069 exons. The merged annotation of all the genes can be browsed and the gene transfer format (GTF) file is freely available to download from the website. Additionally, we have generated a UCSC Genome Browser custom track of the lncRNAKB GTF annotation file that accessible within the website via a html iframe. On the website, users can search the lncRNAKB annotation database by ID, chromosomes, gene start and stop coordinates, gene type, gene names, names of the six lncRNAs sources or any other descriptor and download the search results in .csv format.

### Evaluation of coding potential of lncRNAs

To characterize the lncRNAs annotated in lncRNAKB, FEELnc algorithm was used to classify them based on their position, and their coding potential was evaluated. After applying the FEELnc filters (removing transcripts < 200 bp long and > 75% overlap with protein-coding transcripts, see construction and content of lncRNAKB), the resulting lncRNAKB GTF annotation file included 96,539 genes, 311,241 transcripts and 1,200,236 exons that were considered to be “candidate lncRNAs.” The coding potential score (CPS) cut-off determined by the Random Forest (RF) classification on the training data was 0.434 (separating protein-coding (mRNAs) versus lncRNAs transcripts) with an Area Under the Curve (AUC) performance of 0.972 which maximizes the mRNA classification sensitivity and specificity (see construction and content of lncRNAKB). Based on this cut-off, 83,190 genes, 219,324 transcripts were classified as lncRNAs and 31,402 genes, 91,845 transcripts as protein-coding. The classification module categorized 141,394 lncRNAs transcripts as GENIC (when the lncRNA transcript overlaps an mRNA/protein-coding transcript from the reference annotation file) and 50,540 as INTERGENIC (lincRNAs). Several lncRNA transcripts did not have an interacting mRNA partner thus, remained positionally unclassified. Table 3 summarizes the results of the classifier module with a breakdown of interactions between the two types of lncRNAs and their partner mRNAs/protein-coding transcripts. The lincRNAs are, on average 23kb away from their mRNA partner.

**Table 3:** Summary of classification of lncRNAs transcripts with respect to their localization, overlap and orientation relative to transcription of proximal protein-coding RNA transcripts. The legend below explains the categories in detail: ^1^GENIC: when the lncRNA gene overlaps an RNA gene from the reference annotation file ^2^INTERGENIC (lincRNA): otherwise GENIC type: Then exonic or intronic locations: ^1a^Overlapping subtype: the lncRNA partially overlaps the RNA partner transcript ^1b^Containing subtype: the lncRNA contains the RNA partner transcript ^1c^Nested subtype: the lncRNA is contained in the RNA partner transcript INTERGENIC type: ^2a^Divergent subtype: the lncRNA is transcribed in head to head orientation with RNA partner transcript

| <sup>1</sup>GENIC |  |  |  |  |
| --- | --- | --- | --- | --- |
|  | <sup>1a</sup>Overlapping | <sup>1b</sup>Containing | <sup>1c</sup>Nested | Total |
| Antisense Exonic | 9,326 | 1,816 | 3,552 | 14,694 |
| Antisense Intronic | 1,302 | 1,284 | 8,330 | 10,916 |
| Sense Exonic | 29,942 | 42,160 | 29,087 | 101,189 |
| Sense Intronic | 327 | 994 | 13,274 | 14,595 |
| Total | 40,897 | 46,254 | 54,243 | 141,394 |

| <sup>2</sup>INTERGENIC |  |  |  |  |
| --- | --- | --- | --- | --- |
|  | <sup>2a</sup>Convergent | <sup>2b</sup>Divergent | <sup>2c</sup>Same_Strand | Total |
| Upstream | - | 14,930 | 13,408 | 26,470 |
| Downstream | 11,540 | - | 10,662 | 24,070 |
| Total | 11,540 | 14,930 | 24,070 | 50,540 |

- Then upstream or downstream locations ^2b^Convergent subtype: the lncRNA is oriented in tail to tail with orientation with RNA partner transcript

- Then upstream or downstream locations ^2c^Same_strand subtype: the lncRNA is transcribed in the same orientation with RNA partner transcript

- Then upstream or downstream locations

### Evaluation and comparison of lncRNA and mRNA conservation scores

In addition to the evaluating the coding potential, the conservation of exonic sequences of the lncRNAs and mRNAs was determined (see construction and content of lncRNAKB) and compared. Figure 9 shows the two box plot distributions of exon sequence conservation scores comparing protein-coding and lncRNAs in the lncRNAKB annotation database. Overall, it shows that exons of the protein-coding genes have higher mean sequence conservation scores compared to exons of the lncRNAs. Additionally, the conservation scores for all the exons will be available for review and download at the respective gene page on the website.

### Tissue-specific expression profiling of lncRNAs

To further understand the function of lncRNAs, their gene expression profile across tissues was determined. For this purpose, RNA seq data from 31 solid organ tissues was accessed from GTEx, processed by RNA-seq analysis pipeline, and the gene expression evaluated using the lncRNAKB GTF file (see construction and content of lncRNAKB). Supplementary Table 3 shows the number of RNA-seq samples we analyzed across 31 solid organ human normal tissues from GTEx (*n*=9,425). Supplementary Table 4 shows the summary statistics of alignment (total number of paired-end reads, total number of uniquely aligned paired-end reads, unique and overall alignment rate) across all samples analyzed by tissue. Supplementary Table 5 shows the summary statistics of quantification (total gene count, total number of uniquely aligned paired-end reads used for quantification, total number of uniquely aligned paired-end reads assigned to genes and proportion of successfully assigned paired-end reads to genes) across all RNA-seq samples analyzed by tissue. Supplementary Figure 2 shows the distribution of uniquely aligned paired-end reads assigned to genes across all samples. Bars highlighted in red show the numbers of samples with < 10^6^ reads assigned to genes (*n*=351) that were excluded from further analysis. In the website, users can visualize the normalized gene expression levels (TPM) across 31 solid organ human normal tissues by searching for any gene. Figure 5 shows an example box plot distribution of gene *NPPB* (natriuretic peptide B) for visualization. *NPPB* had a Tau (overall) and PEM score (top five highest positive tissue-specificity score in the heart tissue) of 1 and 1.49 respectively (see: Evaluating tissue-specificity of lncRNAs). It functions as a cardiac hormone and plays a key role in cardiac homeostasis [72]. A high concentration of this protein in the bloodstream is indicative of heart failure. Even though *NPPB* is categorized as a PCG, it has three transcript isoforms that are characterized as lncRNAs. Users can download the boxplot of any gene of interest and download genome-wide gene expression matrices (raw counts and TPM) in text format across all 31 solid organ human normal tissues in the website.

**Figure 5:**
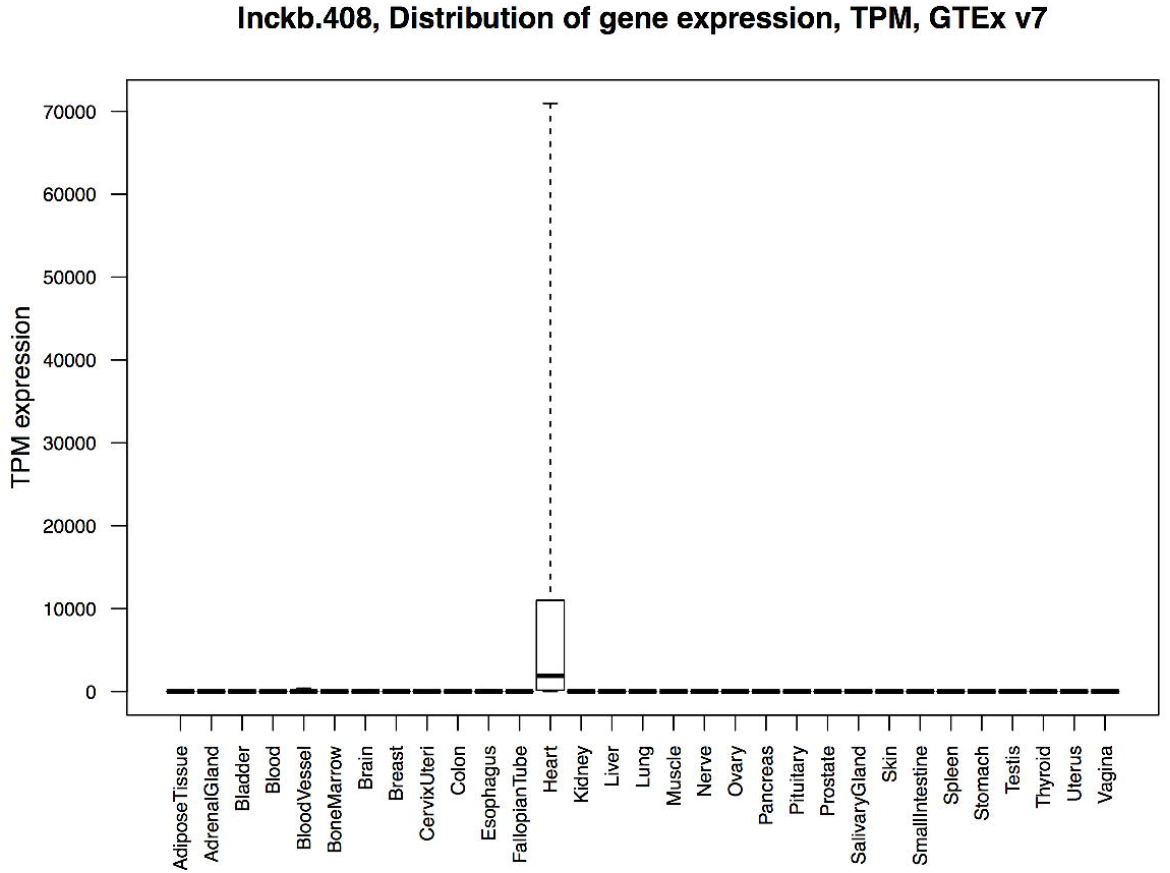
Gene expression box plot distribution of gene *NPPB* (natriuretic peptide B). The x-axis represents the 31 solid organ human normal tissues from GTEx and y-axis is the TPM expression. *NPPB* was ranked among the top five heart-specific genes

### Evaluating tissue-specificity of lncRNAs

Using the gene expression results described in the section above, the tissue specificity score of all the lncRNAs was calculated. Two different metrics, Tau and Preferential Expression Measure (PEM), were calculated that both indicates the tissue specificity of the lncRNAs (see construction and content of lncRNAKB). Figure 6 shows the density distribution of tissue-specificity metrics Tau and PEM across protein-coding genes (PCGs) and lncRNAs in the lncRNAKB annotation database as a comparison. The tissue-specificity scores vary from 0 to 1, where 0 means broadly expressed, and 1 is specific. Figure 6A displays average Tau score across all tissues and Figure 6B displays the maximum and normalized specificity value of PEM among all tissues. Overall, Figure 6 shows that lncRNAs have higher tissue-specificity compared to PCGs. For each gene, we provide a distribution of PEM scores across all tissues and all tissue-specific figures and score files are available for viewing and downloading individually on the lncRNAKB website.

**Figure 6:**
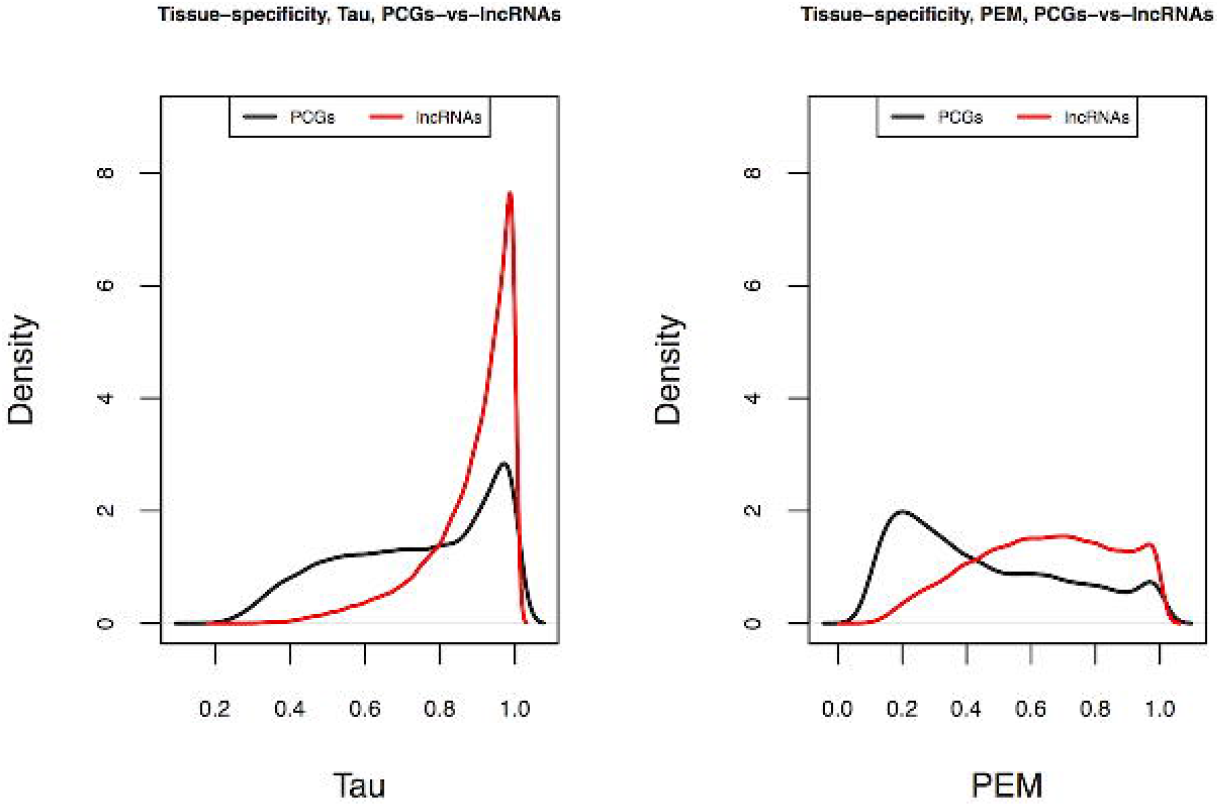
Distribution of tissue-specificity scores with data for RNA-seq of 31 solid organ human normal tissues from GTEx across protein-coding genes (PCGs) and lncRNAs in the lncRNAKB as a comparison. The tissue-specificity scores varies from 0 to 1, where 0 means broadly expressed, and 1 is specific. Graph created with density function from R, which computes kernel density estimates A: Average Tau score across all tissues B: Maximum and normalized specificity value of PEM among all tissues

The tissue specificity of lncRNAs could also be determined by performing an unsupervised principle component analysis (PCA) (see construction and content of lncRNAKB). For this analysis, the log transformed TPM lncRNAs expression data across all tissues was used. Each tissue showed a characteristic transcriptional signature, as revealed by PCA of lncRNA expression. The separation was evident between nonsolid (blood) and solid tissues and, within solid tissues, brain and testis are the most distinct (Fig 7). This finding is an additional confirmation that lncRNAs are tissue-specific.

**Figure 7:**
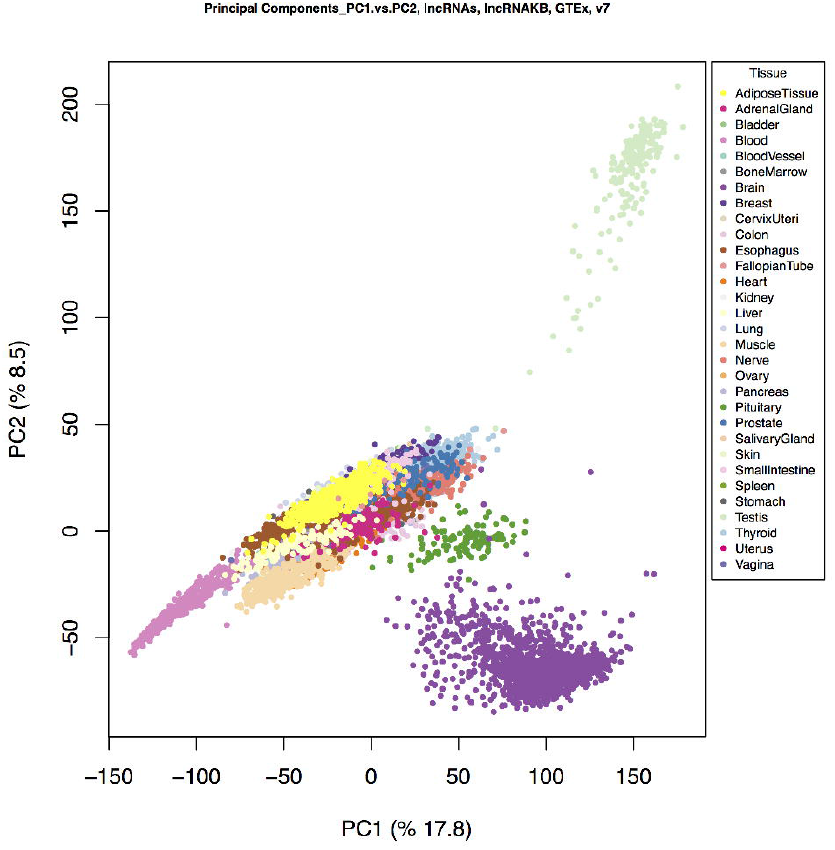
Principal Component Analysis (PCA) of GTEx samples based on lncRNA expression. PCA of all samples based on expression levels of lncRNAs (*log*_2_(*TPM*) transformed). Expression of lncRNAs alone also recapitulates tissue types

### eQTL analysis of lncRNAs

To add to our understanding of lncRNAs gene expression information, we utilized the gene expression data (described in previous sections) in combination with the whole genome sequencing (WGS) data available at GTEx consortium to identify variants in the genome that can alter gene expression (see construction and content of lncRNAKB). This analysis resulted in identification of a number of variants that significantly alter lncRNA gene expression in tissue-specific manner. Table 4 summarizes the results of the *cis*-eQTL analysis. *Cis*-eQTL analysis was performed on 25 tissues that had > 80 samples and also had with WGS. The WGS VCF file with 50,862,464 variants were processed and the resulting file had 5,835,187 SNPs that were used for the *cis*-eQTL analysis (see construction and content of lncRNAKB). For each tissue, Table 4 summarizes the number of samples (stratified by sex), the number of SNPs available after preprocessing, the number of genes that met the TPM threshold criteria from the RNA-seq data (PCG and lncRNAs), the total number of SNP-gene pairs that were tested within 1 Mb of the transcription start site (TSS) of each gene and the number of top *cis*-eQTL genes that met the permutation p-value <= 0.05 threshold after employing the FastQTLv2.0 adaptive permutations approach (see construction and content of lncRNAKB). In the website, users can visualize the *cis*-eQTL results by tissue via a Manhattan plot. Figure 8 shows an example plot from the heart tissue. All figures are available for viewing and download individually on the lncRNAKB website in tissue-specific pages. Users can also download genome-wide compressed *cis*-eQTL results files (text format) by tissue (all SNP-gene pairs and top SNP-gene pairs generated via permutation). Additionally, users can also view and download the SNPs that alter any lncRNA from the gene page.

**Figure 8:**
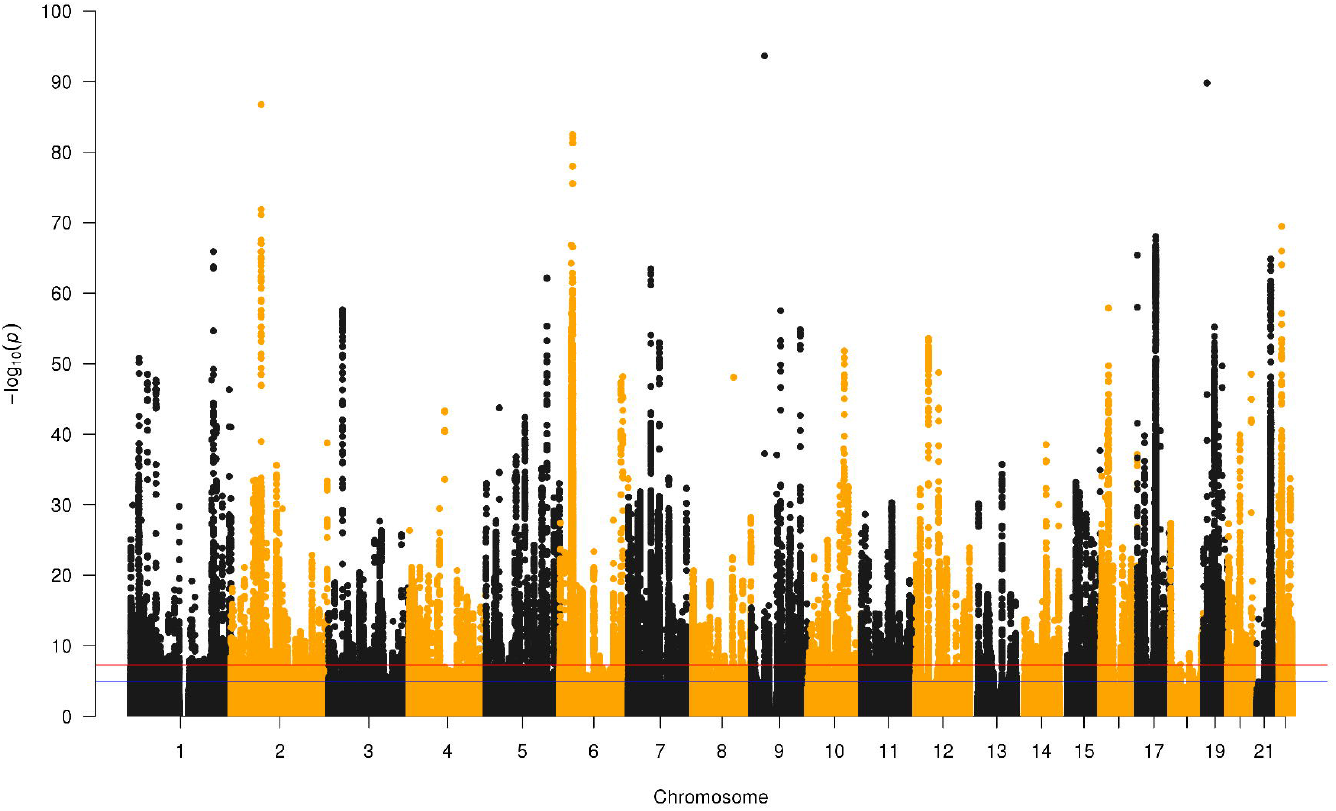
Manhattan plot illustrating the results of the *cis*-eQTL analyses from the heart tissue. The x-axis are the chromosomes and each dot on the y-axis represents the *cis*-eQTL −log10 (p-values) of the SNP-gene pairs that were tested within 1 Mb of the TSS of each gene

**Figure 9:**
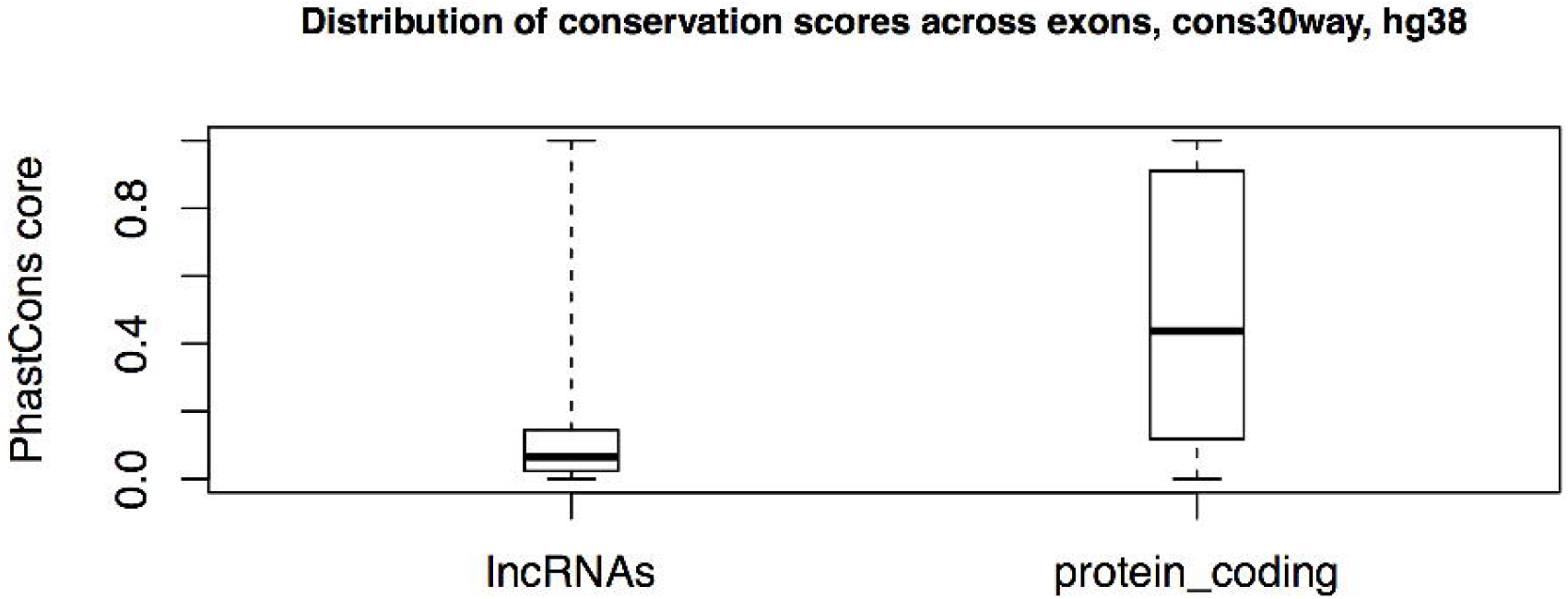
Distribution of mean PhastCons exon sequence conservation scores across lncRNAs and protein-coding genes in the lncRNAKB

**Table 4:** Summary results of the *cis*-eQTL results available from the lncRNAKB. Tissues with < 80 samples are shown here but, were excluded from the analysis *No data due to small sample size or lack of genotype data

| Tissue | Number_of_RNA_seq_samples_with_WGS | Number_of_Males | Number_of_Females | Number_of_SNPs_with_MAF_greater_than_0.05 | Total_number_of_genes_passed_filter | Total_number_of_PCGs | Total_number_of_lncRNAs | Total_SNP_gene_pairs_eQTLs | Total_SNP_gene_pairs_with_permutation_pvalue_less_than_0.05 |
| --- | --- | --- | --- | --- | --- | --- | --- | --- | --- |
| Bladder | 9 | 4 | 5 | 5,462,615 | 28,695 | 15,597 | 13,098 | * | * |
| Blood | 356 | 226 | 130 | 5,953,536 | 18,412 | 11,788 | 6,624 | 37,414,178 | 2,877 |
| Blood_Vessel | 378 | 241 | 137 | 5,963,536 | 25,614 | 14,770 | 10,844 | 51,947,442 | 5,854 |
| Bone_Marrow | * | * | * | * | 22,571 | 12,612 | 9,959 | * | * |
| Brain | 170 | 116 | 54 | 5,857,467 | 31,339 | 16,148 | 15,191 | 62,844,553 | 3,488 |
| Breast | 184 | 102 | 82 | 5,901,708 | 28,839 | 15,680 | 13,159 | 58,130,064 | 4,267 |
| Cervix_Uteri | 8 | 0 | 8 | 5,522,234 | 28,706 | 15,649 | 13,057 | * | * |
| Colon | 250 | 148 | 102 | 5,907,992 | 28,297 | 15,781 | 12,516 | 57,063,773 | 4,767 |
| Esophagus | 353 | 221 | 132 | 5,941,386 | 26,803 | 15,439 | 11,364 | 54,314,052 | 4,815 |
| Fallopian_Tube | 7 | 0 | 7 | * | 18,492 | 16,552 | 1,940 | * | * |
| Heart | 251 | 163 | 88 | 5,913,705 | 24,959 | 14,788 | 10,171 | 50,153,256 | 4,375 |
| Kidney | 29 | 23 | 6 | 5,742,588 | 28,917 | 15,726 | 13,191 | * | * |
| Liver | 118 | 77 | 41 | 5,871,833 | 23,846 | 14,204 | 9,642 | 47,689,780 | 2,759 |
| Lung | 274 | 182 | 92 | 5,926,605 | 29,045 | 15,744 | 13,301 | 58,884,074 | 5,461 |
| Muscle | 359 | 220 | 139 | 5,962,131 | 22,042 | 13,558 | 8,484 | 44,548,539 | 4,454 |
| Nerve | 268 | 174 | 94 | 5,941,274 | 29,326 | 15,472 | 13,854 | 59,363,204 | 7,416 |
| Ovary | 99 | 0 | 99 | 5,873,449 | 27,292 | 14,845 | 12,447 | 54,588,663 | 3,466 |
| Pancreas | 167 | 98 | 69 | 5,905,087 | 23,569 | 14,210 | 9,359 | 47,408,959 | * |
| Pituitary | 108 | 76 | 32 | 5,814,865 | 30,586 | 15,848 | 14,738 | 60,707,019 | 3,949 |
| Prostate | 101 | 0 | 101 | 5,810,666 | 30,373 | 15,931 | 14,442 | 60,377,553 | * |
| Salivary_Gland | 63 | 43 | 20 | 5,771,591 | 28,409 | 15,679 | 12,730 | * | * |
| Skin | 442 | 278 | 164 | 5,966,760 | 27,316 | 15,442 | 11,874 | 55,698,051 | 6,210 |
| Small_Intestine | 90 | 54 | 36 | 5,777,092 | 30,046 | 15,950 | 14,096 | 59,426,622 | 2,987 |
| Spleen | 108 | 62 | 46 | 5,874,443 | 28,284 | 14,969 | 13,315 | 56,914,604 | 4,743 |
| Stomach | 182 | 104 | 78 | 5,890,077 | 26,974 | 15,530 | 11,444 | 54,242,450 | 3,804 |
| Testis | 171 | 0 | 171 | 5,875,543 | 47,909 | 17,777 | 30,132 | 98,376,057 | 8,951 |
| Thyroid | 286 | 183 | 103 | 5,941,584 | 29,715 | 15,604 | 14,111 | 60,217,108 | 7,611 |
| Uterus | 82 | 0 | 82 | 5,795,583 | 28,175 | 15,166 | 13,009 | 55,748,102 | 3,037 |
| Vagina | 87 | 0 | 87 | 5,837,620 | 28,423 | 15,629 | 12,794 | 56,861,978 | 2,865 |

### Functional characterization of lncRNAs using a network-based approach

To further our understanding of lncRNAs, we also undertook WGCNA, a network-based approach that relies on calculating correlation and identifying clusters/modules of correlated between genes (both protein-coding and lncRNAs) (see construction and content of lncRNAKB). Since correlated genes are predicted to play similar functions in the cells, the pathway enrichment analysis of the correlated clusters/modules can help characterize the functions of lncRNAs in the correlated module. Supplementary Table 6 summarizes the results of the WGCNA analysis across the 28 solid organ human normal tissues using the GTEx RNA-seq data. WGCNA analysis was not performed on three tissues due to insufficient sample size. After filtering genes with low expression (see construction and content of lncRNAKB), the average number of protein-coding genes was 14,699 and lncRNAs was 3,389, per tissue. We identified total of 1,208 lncRNA-mRNA co-expression modules across all tissues (on average approximately 43 modules per tissue). On average, across all tissues, each module had approximately 487 genes including 92 lncRNAs, indicating favorable co-expression of lncRNAs with PCGs. Supplementary Table 7 summarizes the results of the over-representation analysis (ORA) based on the hypergeometric test using the Gene Ontology (GO) pathways across all the modules identified in the 31 solid organ human normal tissues. It displays the number of GO pathways tested, number of pathways with p-value ≤ 0.05 and FDR q-value ≤ 0.05 in all modules by tissue. On average, across all modules, each tissue had approximately 2,592 pathways with q-value ≤ 0.05, indicating significant enrichment of biological processes within each of these modules.

Supplementary Table 8 shows the results of WGCNA in heart tissue for all lncRNA-mRNA co-expression modules identified. There were 61 modules identified in the heart using gene expression data across 16,882 protein-coding genes and 2,762 lncRNAs. Supplementary Table 8 separates the number of genes and lncRNAs in each module, representing the size of each. It displays a list of lncRNAs and top 20 hub genes (genes with highest connectivity) in each module. Supplementary Table 9 shows the results of all GO pathways enriched in the heart tissue by module. There were several significant pathways identified (q-value <= 0.05) with many of these involved in heart related biological processes. Figure 10 highlights the network figure created using Cytoscape for module M2 identified in the heart tissue. This module is involved in heart-specific processes such as heart growth, development and contraction. The network has 148 genes (34 protein-coding and 106 lncRNAs) after filtering the adjacency matrix with correlations < 0.20 and “heart development” specific pathways/genes. The orange triangles and green circles/nodes represent lncRNAs and mRNAs respectively. The thickness of the edges highlights the correlation between nodes. The relatively strong connections of several lncRNAs to PCGs in this network suggests these could be potentially involved in the same heart development specific biological processes. On the lncRNAKB website, for each tissue, users can view and download the WGCNA results (similar to Supplementary Table 8) and module enrichment results for all GO pathways (similar to Supplementary Table 9) as comma separated (.csv) files. In addition, 25 notable pathways were selected for each tissues and network files highlighting the lncRNA-mRNA connections were generated. The users can visualize and download the corresponding network figures and review the connections between lncRNAs and mRNAs involved in biological processes of interest.

**Figure 10:**
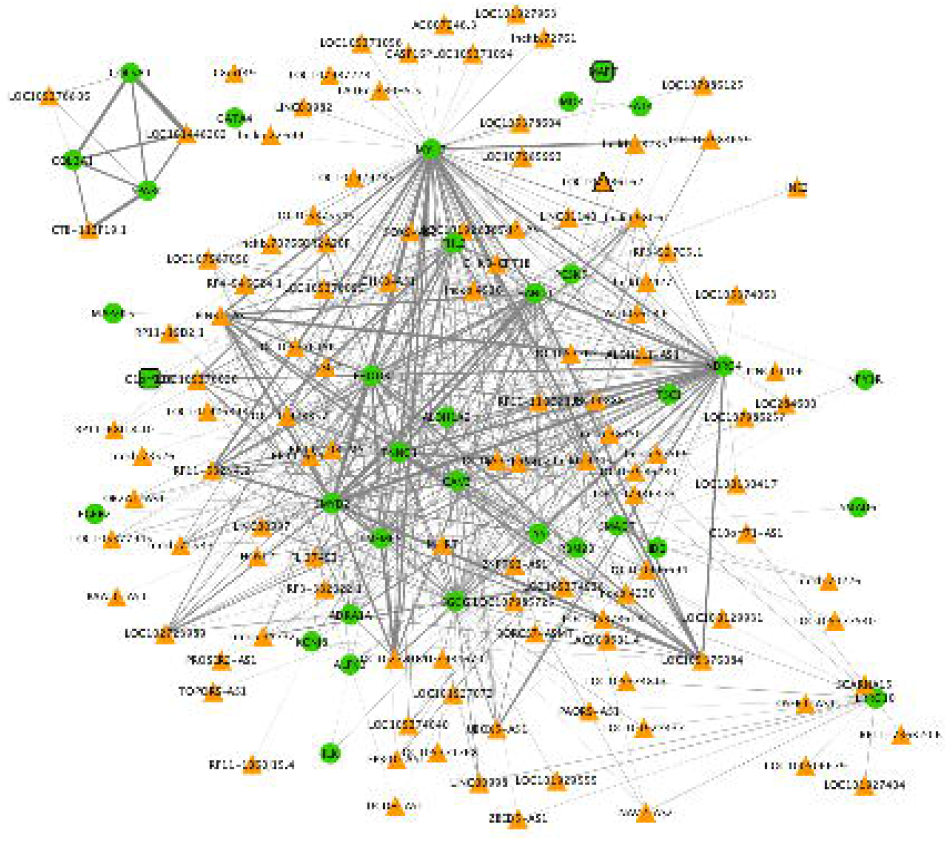
Cytoscape network for lncRNA-mRNA co-expression Module 2 (M2) in the heart identified using WGCNA. The network was filtered for heart development genes (*n*=148) and correlations > 0.20. Orange triangles and green circles/nodes represent lncRNAs and PCGs respectively. The density of gray lines/edges represents the strength of the connection between genes

### Discussion and future directions

There is a large volume of transcriptomics data publicly available and currently being produced at an unprecedented rate. Novel transcripts assembly using RNA-seq data is a method that generates thousands of new transcripts that need to be characterized. Several of these novel transcripts have been categorized as lncRNAs. While these data support the presence of lncRNAs in cell and tissue specific manner, bigger questions surrounding the purpose and functionality of lncRNAs in human biology remain. Several lncRNA databases have tried to address some of these questions, however, there is clearly a need to integrate the lncRNA annotation between databases to create a non-redundant list of well-annotated lncRNA entries that could be utilized by biologists to pursue research in this area.

To address this need, we have created the lncRNAKB, a well-structured research tool that delivers valuable data on human lncRNAs, which can be used for data exploration and hypothesis building purpose. Briefly, the lncRNAKB is the end-product of systematic step-wise integration of six widely used lncRNA databases that resulted in a total non-redundant 99,717 genes entries that were accompanied with 530,947 transcripts and 3,513,069 exon entries. All the annotated lncRNAs can be browsed at http://www.lncrnakb.org. This web-resource also provides a comprehensive list of information that researchers can access for every lncRNA entry. This includes viewing and downloading coding potential, conservation score, tissue specific expression information, and tissue specificity score for any gene of interest from the website. In addition to the gene-level information, we have also created a lncRNA body map where we have utilized the gene expression information and created a tissue specific gene expression and network pages that can be browsed by researchers and the information downloaded as per their research needs. This information includes gene expression count and TPM matrix for all the tissues, eQTL results and lncRNA-mRNA co-expression clusters/modules and the pathway enrichment results of respective modules. Put-together, the above-described features presented in the lncRNAKB web-resource will provide a comprehensive set of information that could be used by biologist interested in pursuing research in lncRNAs in their tissue or biological process of interest.

## CONCLUSION

We have created the <u>l</u>ong <u>n</u>on-<u>c</u>oding <u>RNA</u> <u>k</u>nowledge<u>b</u>ase (lncRNAKB) by methodically integrating widely used lncRNAs resources. We systematically and carefully combined these resources, created a meticulously crafted non-redundant resource and provide valuable functional information on lncRNAs. The present release of lncRNAKB describes a substantial advance in annotations of the largest number of unique lncRNAs (*n*=77,199). In addition, we employed the Genotype-Tissue Expression (GTEx) project to provide tissue-specific expression profiles and tissue-specificity scores for these lncRNAs in 31 solid organ human tissues. We also performed Weighted Gene Co-expression Network Analysis (WGCNA) to identify co-expressed lncRNA-mRNA that were then subjected to pathway enrichment analysis to identify meaningful biological processes that lncRNAs could be potentially involved in, providing potential understanding on lncRNAs function. We created dynamic Cytoscape networks for exploration and visualization of each pathway. The lncRNAKB also incorporates coding potential and phylogenetic conservation. Furthermore, using whole genome sequencing data of 652 subjects from GTEx, we calculated expression quantitative trait loci (*cis*-eQTL) regulated lncRNAs in all tissues. This will provide a strong foundation for integrative traits association analysis by linking a variety of trait related GWAS and our eQTL results. All these features are provided in a user-friendly web interface, available at http://www.lncrnakb.org.

## DECLRATION

### Ethics approval and consent to participate

Not applicable

### Consent for publication

Not applicable

### Availability of data and material

The Genotype-Tissue Expression (GTEx) expression and genotyping data were obtained from the GTEx portal through dbGaP accession phs000424.v7.p2 via available through https://www.ncbi.nlm.nih.gov/projects/gap/cgi-bin/study.cgi?study_id=phs000424.v7.p2

CHESS: http://ccb.jhu.edu/chess/

LNCipedia: https://lncipedia.org/info

NONCODE: http://www.noncode.org/index.php

FANTOM: http://fantom.gsc.riken.jp/5

MiTranscriptome: http://mitranscriptome.org/

BIGTranscriptome: http://bhyou.dothome.co.kr/

All analysis results (raw and normalized tissue specific expression, network modules, and *cis*-eQTL) are available vie lncRNAKB web portal at http://lncrnakb.org under open-source licenses.

### Competing interests

The authors declare that they have no competing interests.

### Funding

This work was supported by NIH grant ZIC-HL006228 to MP.

### Authors’ contributions

FS conceived and designed the experiment, and conducted all aspects of the analysis, generated figures, and drafted the manuscript. KS contributed in database curation, differential expression, WGCNA and eQTL analyses, generated figures and drafting the manuscript. AS, YCC, VC, and IT contributed in database curation, expression and eQTL analysis. XR, PL, YC, and HC contributed in database curation and analysis. RSL, FG, PZ, and MSJ contributed in experimental design and critically evaluated results and manuscript. MP supervised the project, conceived and designed the experiments and analysis, and wrote the manuscript. All authors read and approved the final manuscript.

## Supplementary Figure and Table legends

**Supplementary Figure1:** Illustration showing the different classes of lncRNAs with respect to localization and the direction of transcription of nearby mRNAs (protein-coding genes)

**Supplementary Figure2:** Distribution of successfully assigned RNA-seq reads to lncRNAKB gene annotation. Red bars represent samples with < 10^6^ assigned reads

**Supplementary Table1a:** Number of transcripts and the sources of annotation databases at gene level for protein-coding genes in CHESS

**Supplementary Table1b:** Number of transcripts and the sources of annotation databases at gene level for protein-coding genes in lncRNAKB

**Supplementary Table2a:** Number of transcripts and the sources of annotation databases at gene level for non-coding genes in CHESS

**Supplementary Table2b:** Number of transcripts and the sources of annotation databases at gene level for non-coding genes in lncRNAKB

**Supplementary Table3:** Number of RNA-seq samples that were analyzed across 31 solid organ human normal tissues from GTEx

^1^Excludes samples that did not download (FASTQ files from dbGAP) and/or quantify (via FeatureCounts) successfully

**Supplementary Table4:** Summary statistics of alignment (total number of paired-end reads, total number of uniquely aligned paired-end reads, unique and overall alignment rate) across all RNA-seq samples analyzed by tissue from GTEx

**Supplementary Table5:** Summary statistics of quantification (total gene count, total number of uniquely aligned paired-end reads used for quantification, total number of uniquely aligned paired-end reads assigned to genes and proportion of successfully assigned paired-end reads to genes) across all RNA-seq samples analyzed by tissue from GTEx

**Supplementary Table6:** Summary results of the WGCNA analysis across the 28 solid organ human normal tissues using the GTEx RNA-seq data

**Supplementary Table7:** Summary results of the over-representation analysis (ORA) based on the hypergeometric test using the Gene Ontology (GO) pathways across all the modules identified in the 31 solid organ human normal tissues from GTEx

**Supplementary Table8:** Results of WGCNA in heart tissue for all lncRNA-mRNA coexpression modules identified

**Supplementary Table9:** Results of all GO pathways enriched in the heart tissue by module

