## Supplementary figures and images for "lncRNAKB: A comprehensive knowledgebase of long non-coding RNAs"

### Supplementary Figure1

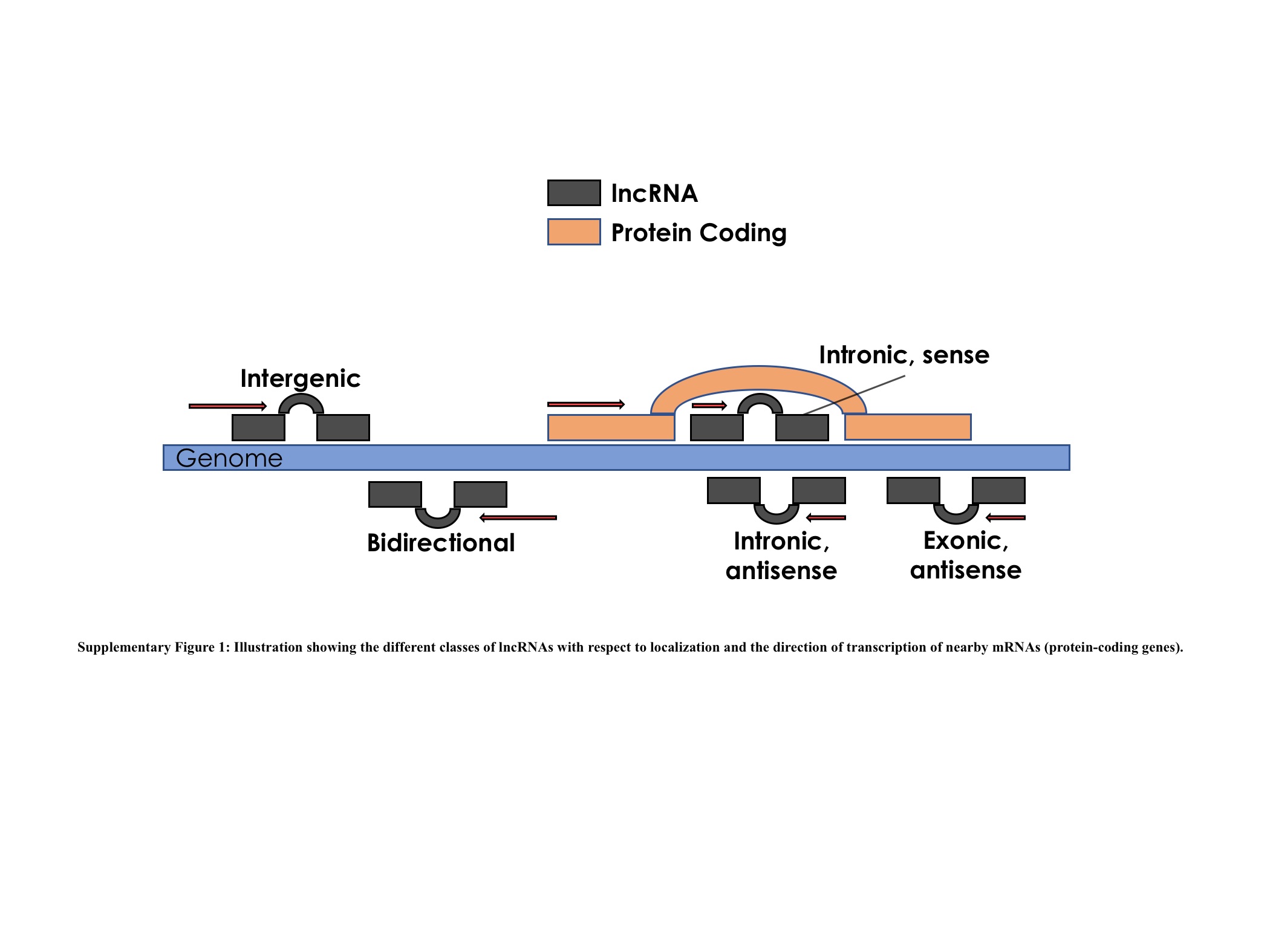

### Supplementary Figure2

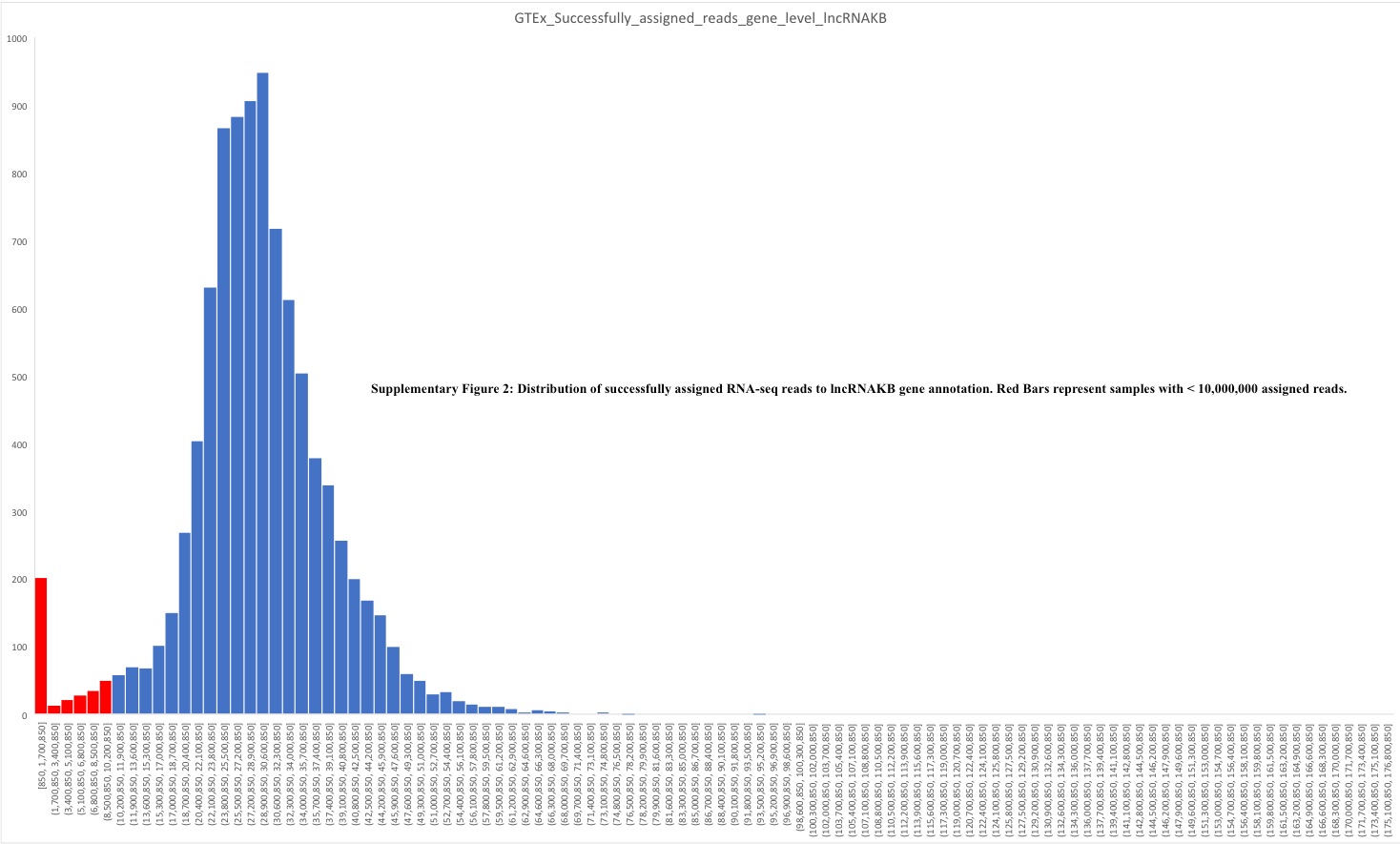
